# Mobbing-like response to secondary predator cues informs group behaviour in wild meerkats

**DOI:** 10.1101/2020.07.02.182436

**Authors:** Isabel Driscoll, Marta Manser, Alex Thornton

**Affiliations:** Centre for Ecology and Conservation, University of Exeter, Penryn Campus, Penryn, Cornwall TR10 9FE, UK; Department of Evolutionary Biology and Environmental Studies, University of Zurich, Winterthurstrasse 190, 8057 Zürich, Switzerland; Kalahari Meerkat Project, Kuruman River Reserve, Northern Cape, South Africa

**Keywords:** animal behaviour, collective decision-making, defensive responses, information transfer, predator cues

## Abstract

The assessment of current risk is essential in informing defensive behaviours. Many animals use cues left behind by predators, known as secondary predator cues (SPCs), to assess risk and respond appropriately. However, meerkats, Suricata suricatta, exhibit seemingly unique mobbing-like responses to these cues. The benefit of this high-intensity recruitment response is unclear, as unlike genuine mobbing, it cannot help to drive the predator away. One potential explanation is that mobbing-like responses promote information gathering and collective decision-making by the whole group. To examine this, we investigated (i) how meerkats’ responses to SPCs differ from mobbing live animals and (ii) the subsequent behavioural changes following a SPC encounter. Using a dataset gathered over a 20-year period, we first compared the rate of SPC recruitment versus the rate of animal mobbing. We then investigated changes in behaviour (alarm calling, sentinel bouts, distance travelled and pup provisioning) in the hour before and after a SPC encounter. Abiotic factors did not affect recruitment rate to SPCs or live animals, or influence the change in behavioural responses following a SPC encounter. The presence of pups reduced response rate to SPCs, but had no effect on animal mobbing rate, supporting experimental findings that responses towards SPCs are unlikely to function as a form of teaching. Alarm calling rate increased and the distance travelled by the group decreased following a SPC encounter, and were unaffected by the presence of pups or abiotic conditions. The results indicate group-level behavioural changes following a SPC encounter, and a greater degree of plasticity in recruitment to SPCs than to live animals. This response plasticity may reflect a context-dependent need to gather information to make collective decisions for defensive behaviour according to the level of threat perceived.

## Introduction

Appropriate defensive responses in the face of predation are vital to survival. However, animals typically face trade-offs between defensive responses and other behaviours, such as foraging (Lima & Dill 1990; Verdolin 2006). Accurately assessing current risk is therefore essential in informing an individual’s or group’s anti-predator responses, minimising the chance of responding inappropriately or unnecessarily. Animals can use a range of cues to inform their defensive behaviours (Lima & Dill 1990; Thorson *et al*. 1998). The direct presence of a predator can be used in assessing risk through visual or acoustic cues, as well as indirect cues of increased risk such as time of day, habitat type, conspecific cues and conspecific body part remains. Secondary predator cues (SPCs), which are cues left in the environment by a predator (Driscoll *et al*. 2020), can also be used in risk assessment. These cues are produced by the predator but are not directly associated with the predator’s current location, and include predator urine, faeces, fur, regurgitation pellets, scent markings and feathers (Driscoll *et al*. 2020). In group-living species, responses to SPCs may play an important role in shaping group-level defensive responses, but this has yet to be investigated.

The use of SPCs can be beneficial in obtaining information about the threat posed without dangerous direct exposure to a predator. For instance, studies have shown that prey species can extract varied information from SPCs, including the type of predator (Van Buskirk 2001; McGregor *et al*. 2002; Mella *et al*. 2014), its size (Kusch *et al*. 2004), predator density and proximity (Ferrari *et al*. 2006), diet (Mathis & Smith 1993; Apfelbach *et al*. 2015) and how recently the predator may have been in the area (Barnes *et al*. 2002; Zöttl *et al*. 2013; Kuijper *et al*. 2014; Van Buskirk *et al*. 2014). This information can then guide an array of behavioural defensive responses. The most common responses are cue avoidance (of either the cue itself (Caine & Weldon 1989) and/or the area the cue was encountered (Grostal & Dicke 1999; Sike & Rózsa 2006; Amo *et al*. 2011; Weiss *et al*. 2015)) and increased vigilance (e.g. Monclús *et al*. 2005; Zöttl *et al*. 2013; Garvey *et al*. 2016; Tanis *et al*. 2018).

In some animals, initial responses to SPCs involve protracted inspection, presumably as a means of gathering further information (Belton *et al*. 2007; Furrer & Manser 2009; Zöttl *et al*. 2013; Mella *et al*. 2014; Garvey *et al*. 2016). Several social mongoose species, including dwarf mogooses, *Helogale parvula*, banded mongooses, *Mungos mungo*, and meerkats, *Suricata suricatta*, also produce distinctive recruitment calls to gather group members around the cue (Furrer & Manser 2009; Zöttl *et al*. 2013; Collier *et al*. 2017). Meerkats seem to take this a step further by responding in a similar way to encountering a live predator (for photos see: Driscoll *et al*. 2020). This intense, mobbing-like response, involving tail raising, piloerection and recruitment calls, is a seemingly disproportionate response to something that in itself does not pose a threat. The function of this overt reaction is unclear, as it does not provide the primary benefit that mobbing of an actual predator does: driving a threat away (Curio *et al*. 1978; Graw & Manser 2007). Furthermore, experimental evidence shows that exaggerated responses to SPCs are not used as a means of teaching naïve pups about potential threats, since adults decrease their response intensity in the presence of pups (Driscoll *et al*. 2020).

Whereas previous studies have typically focused on individual responses to SPCs, it is possible that meerkats’ unusual mobbing-like response to SPCs may instead serve to inform and influence subsequent group-level behaviour. To maintain one of the primary benefits of group living, reduced risk of predation (Krause & Ruxton 2002; Caro 2005), it may be necessary for all group members to be informed of current risks. Recruitment to SPCs may allow group members to gather detailed information about the threat through inspection, using this information to adjust vigilance behaviours, and aid in increasing group cohesion by bringing group members to a focal point. The exaggerated mobbing-like response may increase likelihood to recruit by denoting a high-level threat and provide a clear signal to move towards. Previous work has shown that meerkats increase individual vigilance during and immediately following an experimental SPC encounter (Zöttl *et al*. 2013). However, this work primarily focused on immediate behavioural changes (minutes after interaction) and predator model detection and not more prolonged changes to group behaviour in the following hour. The effects of encountering a SPC may lead to other changes in behaviour, not yet examined, that may improve our understanding of the use and importance of SPCs in informing defensive behaviours.

Meerkats are cooperative breeders living in the arid regions of southern Africa, in groups ranging from 3-47, averaging 15 individuals (Clutton-Brock & Manser 2016). They forage as a cohesive group, using vocalisations to alert others about potential threats (Manser 2001; Manser *et al*. 2001), maintain proximity between individuals (Gall & Manser 2017) and make collective decisions to move from one foraging patch to the other based on quorum decisions (Bousquet *et al*. 2011). During foraging there is often a sentinel on guard, undertaking vigilance for the whole group, allowing other group members to reduce vigilance and maximise foraging (Clutton-Brock *et al*. 1999; Manser 1999; Santema & Clutton-Brock 2013; Rauber & Manser 2017). Pups begin foraging with the group at around 20-25 days old (Clutton-Brock *et al*. 2000). All group members contribute to offspring care, provisioning pups with food until around three months old (Clutton-Brock & Manser 2016) and teaching pups to hunt by progressively introducing them to live prey (Thornton & McAuliffe 2006). Meerkats mob a variety of threats, primarily predators such as snakes, wildcats and other mammalian predators, but also occasionally non-predators such as hares and antelope (see Graw & Manser 2007 for detailed list). Meerkats also show a mobbing-like response to scents and objects such as fur, predator faeces or urine, owl pellets and feathers (Zöttl et al., 2013, MS1).

This study investigates the potential function of the mobbing-like response in facilitating group-level behavioural changes following natural SPC encounters. Using long-term observational records, we begin by investigating the social and abiotic factors influencing the rate of recruitment to SPCs compared to that of live animals, to see where the differences lie between these almost identical behaviours. We predicted that live animal mobbing events would be less affected by group composition and environmental conditions due to the necessity of responding to an imminent threat. We then investigate group behavioural changes following a SPC encounter, examining alarm calling rate, sentinel rate, distance travelled by the group and pup provisioning rate in the hour prior to and post a SPC recruitment event. We predicted that if responding to SPCs functioned in initiating group defensive behaviours, (1) alarm calling rate and (2) sentinel behaviour, as a cooperative form of vigilance, would increase following a SPC encounter due to an increase in perceived risk and threat sensitivity. We also predicted that (3) the per hour distance travelled would increase following a SPC recruitment event to move out of an area of higher risk and, (4) pup provisioning rate would decrease as a result of a trade-off with defensive responses. In addition, we examined the effect of group composition and group size on these behavioural measures and the effect of current climatic conditions, to investigate response plasticity and variation with social and abiotic conditions.

## Methods

### Study site & population

In this study we analysed 20 years of behavioural data collected as part of long-term monitoring of the meerkat population at the Kalahari Meerkat Project, Kuruman River Reserve (26°59’ S, 21°50’ E) in South Africa (Clutton-Brock *et al*. 1998). For detail about habitat and climate see (Russell *et al*. 2002). All individuals were habituated to human observation (< 1m) and identifiable from unique dye mark patterns on their backs (Jordan *et al*. 2007). Life history was known for the majority of individuals from birth, including age, sex and dominance status, with the exception of immigrating individuals. We analysed data from 11/04/1999 to 30/04/2019, using only observations with complete records for each analysis.

### Data collection

As part of long-term monitoring, meerkat groups were visited at least every three days for a minimum of one hour in either the morning following the group leaving the sleeping burrow and/or the evening prior to the group’s return to the sleeping burrow. During sessions behavioural data was recorded *ad libitum* (every time a behaviour occurs; (Altman 1974)). For definitions of behaviours and other data recorded and analysed as part of this study see Table 1.

**Table 1 –.** Descriptions of data recorded as part of long-term monitoring of the population and analysed as part of this study.

| Data recorded | Description |
| --- | --- |
| Alarm events | When >50% of the group respond to a potential threat. Recording the response given by the majority of the group, responses included: look briefly, watch continuously, move, move to bolthole, move below ground and recruit. Also recorded the type of threat responded to. |
| **Sentinel bouts** | *A period of vigilance undertaken by an individual of over 10 seconds from a vantage point, which an individual has gone out of its way to guard from or is making sentinel calls.* |
| **Pup provisioning** | *An individual (non-pup) gives a pup a food item.* |
| **GPS location** | *GPS fixes taken from the centre of the group at approximately 15 minute intervals during a session (accuracy: 95% of fixes within 5 m; eTrex H, Garmin International Inc., Olathe, KS, USA). Taken either from the time a group left the sleeping burrow or until the group returned to the sleeping burrow.* |
| **Group size and composition** | *The number of individuals present in a session, their sex and age class/dominance categories.* |
| **Daily maximum temperature (°C)** | *Daily maximum temperature* |
| **Daily rainfall (mm)** | *Daily total rainfall* |

Recruitment events were recorded as alarm events (Table 1) and defined as when the group recruits to a stimulus with erect fur and tails, making recruitment calls, often spitting and growling (Manser *et al*. 2014). The eliciting stimulus was also recorded with the animal type or as ‘scents’ or ‘objects’. If the type of threat responded to was not known, it was recorded as ‘unknown’ and excluded from our analyses.

The daily maximum temperature and the total rainfall over the previous 30 days were used as indicators of current habitat conditions and food abundance (Thornton 2008; Hodge *et al*. 2009; Wiley & Ridley 2016), while the total rainfall over the previous 9 months served as an indicator of the groups’ overall condition, in line with previous literature (English *et al*. 2012).

### Data analysis

#### (1) Recruitment event rate (to SPCs or live animals)

Recruitment event rate to SPCs (scents and objects) and live animals (any predator or non-predator) was calculated per group. We analysed data from 54 groups over 20 years, comprising a total of 3705 SPC recruitment events and 1545 mobbing events over 131,289 hours of recorded data over 52,776 sessions. Recruitment rate was calculated over a month period by dividing the number of recruitment events by the number of hours of adlib data recorded, to control for sampling effort.

#### (2) Behavioural changes following a SPC recruitment event

To examine behavioural changes following a SPC recruitment event, we compared the total hourly number of alarm events (not including recruitment events), number of sentinel bouts, distance travelled by the group and per pup provisioning rate, in the hour before and after a SPC recruitment event. The total number of alarm events to potential threats (as outlined above), in an hour was used to indicate changes in threat perception. The total number of sentinel bouts in an hour period was used to indicate changes in cooperative vigilance and perceived risk. The distance travelled in an hour, calculated from the GPS fixes, was used to determine changes in the rate of movement following a recruitment event. Hourly per pup provisioning rate, the number of pup feed events recorded divided by the number of pups present in the group, was used as an indicator to assess the effect of a recruitment event on the maintenance of pup care.

### Statistical analysis

Statistical analysis was conducted using R version 1.1.463 (R Core Team 2015), with packages *lme4* and *glmmTMB* for mixed models. Model assumptions were checked using residual plot distribution techniques. An information theoretic (IT) approach was applied for model selection, using Akaike’s information criterion (AIC) to rank the models following the approach used by Richards *et al*. (2011). Model building was conducted using combinations of fixed effects defined *a priori*. Models within AIC ≤ 6 of the model with the lowest AIC value formed the ‘top set’. To avoid retaining overly complex models we applied the ‘nesting rule’, removing more complex versions of nested models from the top set.

### Recruitment event rate

To analyse the factors influencing rates of recruitment in response to SPCs (model a) and live animals (model b) we used linear mixed models (LMMs). Recruitment rate, using the normalised total number of session hours, was square-root transformed to meet model assumptions of normal data distribution. To understand the influence of social factors, both models (a and b) included group size (the average number of individuals in the group during the month), average sex ratio (mean number of females:males during the month) and the average proportion of pups (proportion of the foraging group that were pups) as explanatory terms. We also included whether or not pups were foraging with the group or not as a categorical explanatory factor to test whether pup presence alone was enough to influence recruitment frequency. Average daily maximum temperature and total rainfall for the previous 30 days, and total rainfall for the previous nine months, were used to test the effect of abiotic conditions on recruitment event frequency. The group identity nested within year was included as a random term to account for repeated measures within groups and variation in group characteristics over the study period.

### Behavioural changes following a SPC recruitment event

Generalised linear mixed models (GLMMs) were used to analyse behavioural changes before and after a SPC recruitment event, examining the hourly number of alarm events (c), hourly number of sentinel bouts (d), metres travelled per hour (e) and hourly provisioning rate per pup (f). Models (c) to (e) were fitted with a negative binomial error structure to account for overdispersion of data, and model (f) was fitted with a zero-inflated negative binomial error structure to account for zero-inflation and overdispersion. The same explanatory terms outlined above were used except sex ratio. Sex ratio was initially included in the analyses but never ranked within the top 5 models with the lowest AIC values during model selection, so was removed to allow more complete records to be analysed. The group size, proportion of pups and daily maximum temperature recorded at the time of the recruitment event were used rather than the monthly average. Model (f) did not include whether pups were foraging with the group as pup provisioning would only occur if they were present. Models (c-f) also included whether the response was in the hour before or after the mobbing to test whether there was a behavioural change, and whether the cue type was a scent or an object to examine whether cue type influences changes in behaviour. The interactions of both before/after and the presence of pups with all other fixed effects were used as both these factors may have interacted with the other factors to influence behaviour. The interaction of average daily temperature with either 30 day and nine-month rainfall was included as temperature and rainfall-driven food availability and body condition are generally closely linked (English *et al*. 2012). Random terms of group nested within year were also included.

## Results

### Recruitment event rate

#### (a) SPC recruitment rate

The interaction between group size and proportion of the group composed of pups influenced the frequency of SPC recruitment events, as did the presence of pups alone. The hourly mobbing rate for each group over a one-month period ranged from 0 to 0.28 (mean±SE = 0.03±0.00), with a total of 3705 SPC recruitment events recorded. LMM analyses produced two models in the top set, both of which were retained following the application of the nesting rule (model a13 and model a12; Appendix Table 2). Model a13 contained the proportion of pups, the average group size for that month and the interaction between the two (estimate(SE) = −0.06(0.04), *χ^2^* = 10.75, d.f. = 1, p < 0.001; Appendix Table 1). When no pups were present in the group, the recruitment rate increased with group size, but as the proportion of pups in the group increased the effect of group size lessened, with a lower recruitment rate with an increasing proportion of pups (Figure 1A). Model a12 was similar, containing whether or not pups were foraging with the group, the average group size and the interaction between the two. SPC mobbing rate was reduced when pups were foraging with the group, although the effect size was rather small (estimate(SE) = −0.29(0.15), *χ^2^* = 22.95, d.f. = 1, p < 0.001; Figure 1B; Appendix Table 1). There was no interaction between the presence of pups and group size (pups foraging*group size: estimate(SE) = −0.001(0.008), *χ^2^* = 0.01, d.f. = 1, p = 0.94; Appendix Table 1). Abiotic factors of rainfall and temperature did not appear in the top set.

**Figure 1 –.**
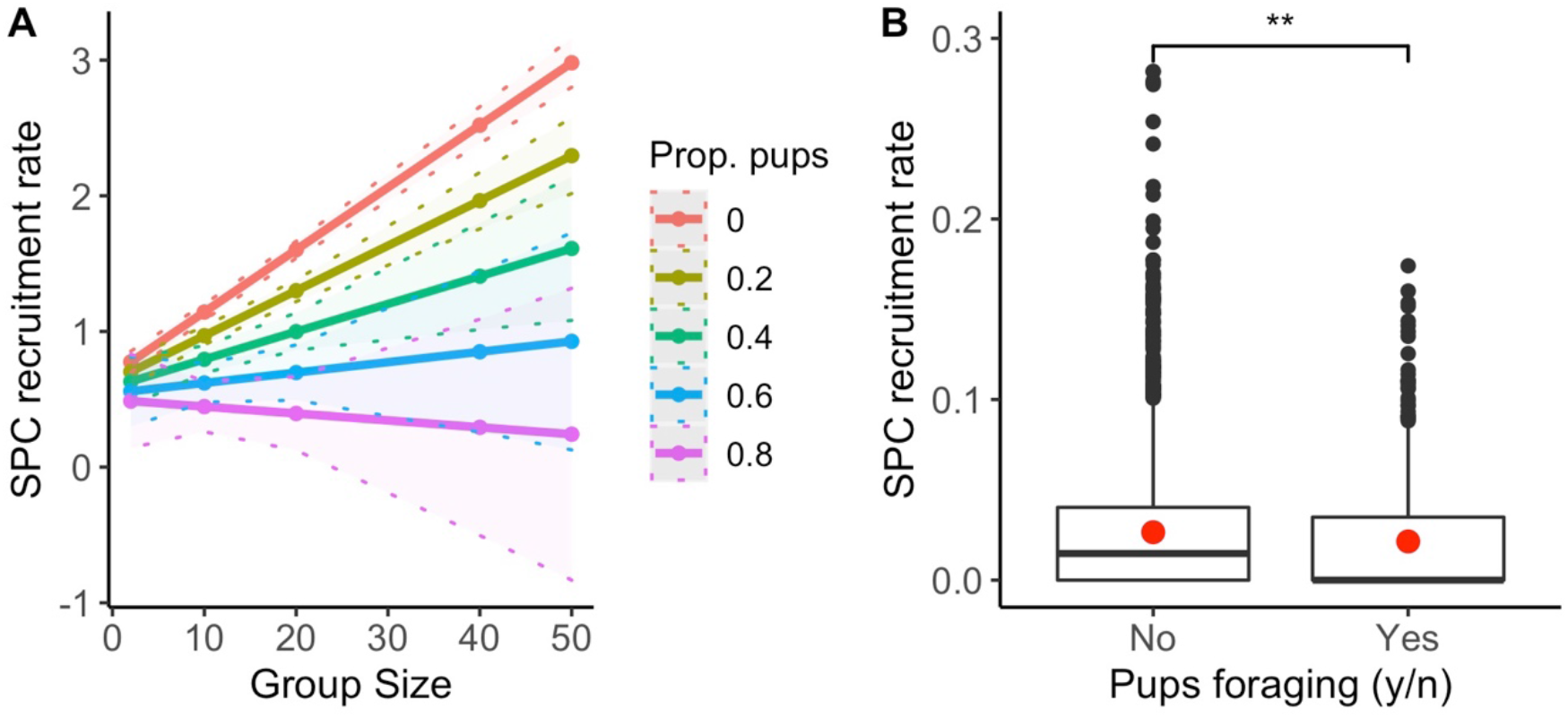
The hourly rate of recruitment events per group in a month to SPCs (months analysed n = 2918) predicted by (A) the interaction between group size and the proportion of the group made up by pups, and (B) whether pups were foraging with the group (yes (n = 660) or no (n = 2258)). The linear regression lines with the shaded area in (A) illustrate the 95% confidence interval. Red dots in (B) indicate the mean rate of recruitment events (yes = 0.02, no = 0.03). Asterix indicating significance level “***” = p < 0.001, “**” = p < 0.01, “*” = p< 0 (B).

#### (b) Animal mobbing rate

The frequency of live animal recruitment events was influenced by group size. Meerkat groups mobbed animals between 0 and 0.26 times per hour each month (mean±SE = 0.01±0.00), with a total of 1545 mobbing events recorded. LMM analyses produced three models in the top set, of which one (model b6; Appendix Table 3) was retained following the application of the nesting rule. Model b6 contained only the average group size as a significant positive predictor of animal mobbing rate (LMM: estimate(SE) = 0.03(0.003), *χ^2^* = 68.59, d.f. = 1, p < 0.001; Figure 2; Appendix Table 1). The proportion of pups as a negative predictor and a slight reduction in mobbing frequency when pups were foraging with the group also appeared in the top set, but did not have a robust effect (proportion of pups: estimate(SE) = −0.22(0.39), *χ^2^* = 2.83, d.f. = 1, p = 0.09; pups foraging: estimate(SE) = −0.04(0.11), *χ^2^* = 0.61, d.f. = 1, p = 0.44; Appendix Table 1), and did not interact with average group size (group size*proportion of pups: estimate(SE) = −0.001(0.03), *χ^2^* = 0.03, d.f. = 1, p = 0.87; group size*pups foraging: estimate(SE) = 0.001(0.01), *χ^2^* = 0.001, d.f. = 1, p = 0.94; Appendix Table 1). Abiotic factors of rainfall and temperature did not appear in the top set.

**Figure 2 –.**
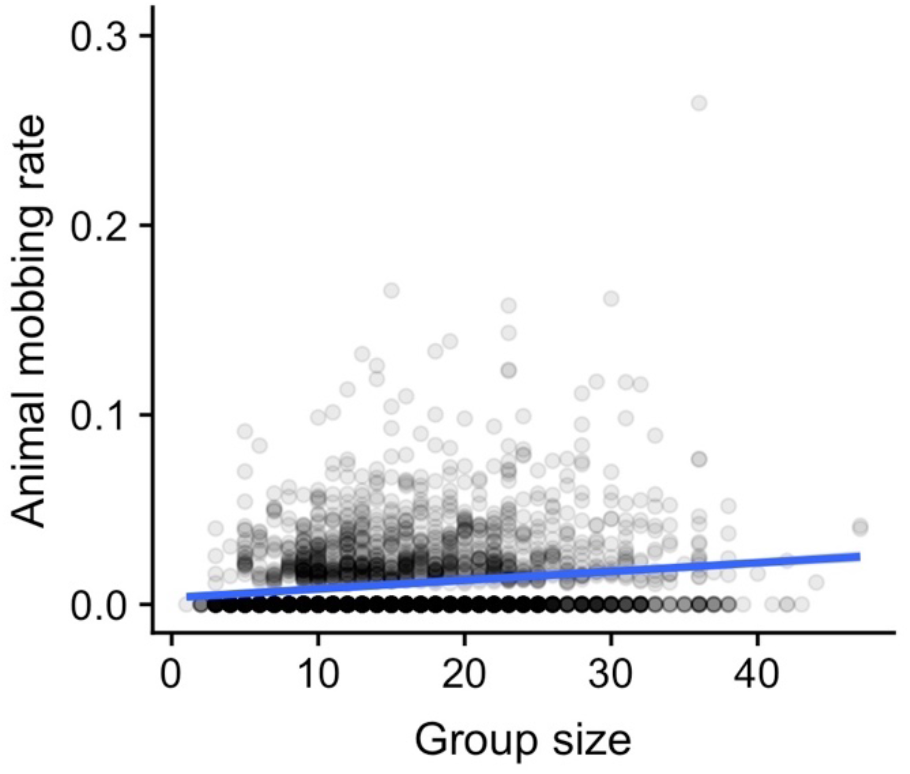
The hourly per group rate of recruitment events in a month to animals predicted by total group size (months analysed n = 2918). Shade of points indicating frequency of data points overlapping. Blue linear regression line with the shaded area illustrating the 95% confidence interval.

### Behavioural changes following a SPC recruitment event

#### (c) Number of alarm calls

We found evidence that the rate of alarm calling increased in the hour following SPC recruitment events and was further influenced by abiotic conditions. The number of alarm calls ranged from 0-19 per hour (mean± SE = 2.23±0.03). GLMM analyses produced two models in the top set of which both (model c8 and c12: Appendix 2 Table 5) were retained following the application of the nesting rule. Model c8 contained the hour before and after the recruitment event, the maximum temperature on that day and the interaction between the two. Alarm calling rate increased in the hour after a recruitment event (2.32±0.04), compared with the hour before (2.14±0.04) (GLMM: estimate(SE) = 0.03(0.11), *χ^2^* = 13.34, d.f. = 1, p < 0.001; Figure 3A; Appendix Table 3), but declined as the daily maximum temperature increased (estimate(SE) = −0.02(0.003), *χ^2^* = 73.61, d.f. = 1, p < 0.001; Figure 3B; Appendix 2 Table 3). There was no interaction between the time period (before/after) and the daily maximum temperature (before/after*temperature: estimate(SE) = −0.004(0.004), *χ^2^* = 1.22, d.f. = 1, p = 0.27; Appendix 2 Table 3). Model c12 contained daily maximum temperature, total rainfall over the last 9 months and the interaction between the two (temperature*rainfall: estimate(SE) = 0.05(0.02), *χ^2^* = 8.09, d.f. = 1, p = 0.004; Appendix 2 Table 3). When rainfall had been high in the previous nine months, the daily maximum temperature had little effect on alarm rate. As nine-month rainfall decreased the disparity between alarm calling rate increased, with alarm calling rate being greater at low temperatures and reduced at high temperatures.

**Figure 3 –.**
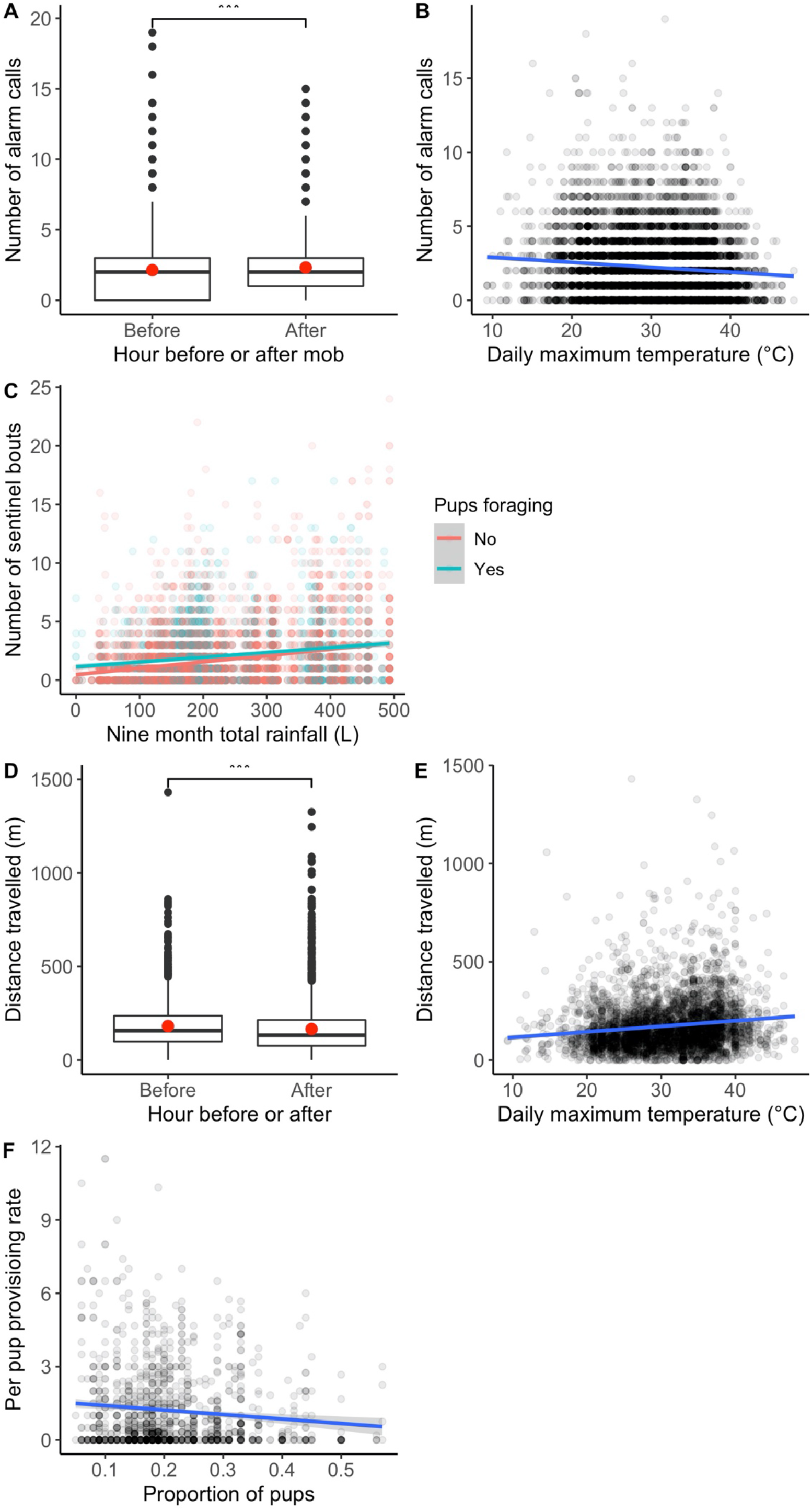
(A) The influence of whether it was before (n = 3473) or after (n = 3473) a SPC recruitment on event hourly alarm calling rate. (B) The influence daily maximum temperature on hourly alarm calling rate (n = 6946). (C) The influence of total rainfall over the last 9 months interacting with whether pups were foraging with the group yes (blue; n = 1806) or no (pink; n = 5138) on hourly number of sentinel bouts (total n = 6944). (D) The influence of whether it was the hour before (n = 2012) or after (n = 2012) a SPC recruitment event on hourly distance travelled by a group. (E) The influence of predicted by daily maximum temperature on hourly distance travelled by a group (n = 4024). (F) The influence of the proportion of the foraging group made up by pups on hourly per pup provisioning rate (n = 1248). Red dots (A & D) indicating mean rate of recruitment events. Shade of points in (B, C, E & F) indicating frequency of data points overlapping. Blue linear regression line for (B, E & F) and blue or pink for (C) with the shaded area illustrating the 95% confidence interval. Asterix indicating significance level “***” = p < 0.001, “**” = p < 0.01, “*” = p< 0.05 (A & D).

#### (d) Number of sentinel bouts

We found evidence that the frequency of sentinel bouts did not change in the hour following a SPC recruitment event, but was increased generally when pups were present and with increasing nine-month rainfall. The number of sentinel bouts ranged from 0-24 per hour (mean± SE = 1.80±0.03). There was no evidence that meerkats increased their rate of sentinel bouts in the hour after encountering a SPC. Instead, GLMM analyses produced one model in the top set following application of the nesting rule (model d19; Appendix Table 6). Model d19 contained whether pups were foraging with the group, rainfall for the previous nine months and the interaction between the two (9 month rainfall*pups foraging: estimate(SE) = − 0.85(0.33), *χ^2^* = 6.50, d.f. = 1, p = 0.011; Figure 3C; Appendix Table 3). Sentinel rate increased with greater rainfall over the previous nine months. Overall, sentinel rate was higher when pups were foraging with the group. Sentinel rate was lower when pups were not foraging with the group, particularly at low rainfall, however, there was little difference between when pups were or were not present at high rainfall.

#### (e) Distance travelled

We found evidence that the hourly distance travelled by meerkat groups reduced in the hour following a SPC recruitment event and was influenced by abiotic conditions. The distance that meerkat groups travelled ranged from 0-1431m per hour observation (mean± SE = 173.59± 2.10 m), using GPS fixes taken at ~15min intervals. GLMM analyses produced three models in the top set, of which all three (model e8, model e12 and model e13; Appendix Table 7) were retained following the application of the nesting rule. Model e8 contained the hour before or after the mobbing event, daily maximum temperature and the interaction between the two. Meerkats travelled shorter distances in the hour following an SPC encounter than the hour before (GLMM: estimate(SE) = 0.27 (0.17), *χ^2^* = 18.23, d.f. = 1, p < 0.001; Figure 3D; Appendix Table 3), and travelled further as daily maximum temperature increased (estimate(SE) = 0.02(0.003), *χ^2^* = 68.11, d.f. = 1, p < 0.001; Figure 4E; Appendix Table 3). The interaction between hour before or after and daily maximum temperature was not robust (before/after*temperature: estimate(SE) = −0.01(0.004), *χ^2^* = 2.12, d.f. = 1, p = 0.15; Appendix Table 3). Model e12 contained daily maximum temperature, total rainfall in the previous nine months and the interaction between the two. Distance travelled reduced as nine month total rainfall increased (estimate(SE) = 0.34(0.69), *χ^2^* = 13.10, d.f. = 1, p < 0.001; Appendix Table 3). The interaction between nine month rainfall and maximum daily temperature did not have a robust effect (9 month rainfall*temperature: estimate(SE) = −0.03(0.02), *χ^2^* = 2.47, d.f. = 1, p = 0.12; Appendix Table 3). Model e13 contained daily maximum temperature, total rainfall for the previous 30 days and the interaction between the two. There was an interaction between 30 days rainfall and daily maximum temperature (30 day rainfall*temperature: estimate(SE) = −0.17(0.08), *χ^2^* = 4.48, d.f. = 1, p = 0.03; Appendix Table 3). At low 30-day rainfall distance travelled increased with temperature, at intermediate rainfall there was a slight increase in distance travelled at higher temperatures. In contrast, at very high rainfall levels, the distance travelled decreased with increasing temperature.

#### (f) Provisioning rate

We found evidence that the hourly per pup provisioning rate did not change in the hour following a SPC recruitment event and was influenced only by the proportion of pups present in the group. The hourly provisioning rate per pup ranged from 0 to 11.5 (mean± SE = 1.20±0.05). GLMM analyses produced three models in the top set, of which one was retained following application of the nesting rule (model f15; Appendix Table 8). Model f15 contained only the proportion of the pups present with per pup provisioning rate decreasing as the proportion of pups increased (GLMM: estimate(SE) = −2.86(0.63), *χ^2^* = 22.52, d.f. = 1, p < 0.001; Figure 3E; Appendix Table 3). Total group size and the hour before or after the mob also appeared in the top set but neither had a robust effect (group size: estimate(SE) = 0.01(0.02), *χ^2^* = 1.25, d.f. = 1, p = 0.26; before/after: estimate(SE) = 0.07(0.17), *χ^2^* = 0.23, d.f. = 1, p = 0.63; Appendix Table 3), and neither interacted with the proportion of pups (group size*proportion of pups: estimate(SE) = −0.15(0.08), *χ^2^* = 3.61, d.f. = 1, p = 0.06; before/after*proportion of pups: estimate(SE) = −0.15(0.76), *χ^2^* = 0.04, d.f. = 1, p = 0.84; Appendix Table 3).

## Discussion

Secondary predator cues (SPCs) can provide information about likely threat levels, but the function of meerkats’ highly exaggerated, mobbing-like response is unclear. We examined whether the information gathered during the active recruitment to SPCs is used to inform subsequent defensive behaviours. Following a SPC encounter alarm calling rate increased and distance travelled decreased, suggesting SPC interactions influence group-level defensive behaviours and collective-decisions. Additionally, there appeared to be a greater degree of plasticity in responding to SPCs, with a larger effect of social conditions on rate of SPC recruitment events compared to live animal mobbing events.

Our results support the idea that the information gained during mobbing-like responses towards SPCs is used to inform subsequent group behaviour. Both the rate of responding to SPCs and live animal mobbing was associated positively with group size. A primary cause of this increased predator detection in larger groups may be due to more potential detectors (Krause & Ruxton 2002; Caro 2005). For example, larger groups of chestnut-crowned babblers, *Pomatostomus ruficeps*, are more likely to encounter predators but less likely to be attacked (Sorato *et al*. 2012). In defending against predators larger groups have greater success and individuals mob with greater intensity (Krams *et al*. 2009). As well as the safety in numbers effect (Hamilton 1971; Lehtonen & Jaatinen 2016), recruiting individuals may act in informing them of the nature of the threat to better inform the group’s collective defensive behaviours. The greater frequency of recruitment events in larger groups, particularly for SPCs, may also be due to difficulties in maintaining group cohesion in larger groups (Focardi & Pecchioli 2005). Recruitment to cues may bring the group to a focal point facilitating informed decision making and cohesive movement away, possibly through quorum sensing. White-breasted mesites, *Mesitornis variegatus*, for example, increase group cohesion following an alarm event (Gamero & Kappeler 2015), and convict cichlids, *Amatitlania nigrofasciata*, do so in response to conspecific alarm cues (Brown & Foam 2004). Whether SPC encounters do result in increased group cohesion in meerkats could be examined by experimentally investigating whether inter-individual distance and the spread of the whole group changes following a recruitment event.

Further evidence suggesting that the mobbing-like response to SPCs may help to inform subsequent group behaviour comes from the increase in alarm calling rate and decrease in distance travelled following an SPC encounter. The increased alarm calling rate may be suggestive of either an increase in actual risk, i.e. a predator in the vicinity, or increased sensitivity to potential threats. Previous work has shown that predator detection latency in meerkats reduces following an SPC encounter (Zöttl *et al*. 2013), indicating that SPCs may be a reliable indicator of a predator in close-proximity and aid in locating the threat. Increased perceived risk may also increase threat-sensitivity resulting in increased alarm calling, as predicted by signal-detection theory (Wiley 2006; Ferrari *et al*. 2009). Under high perceived risk it may be safer to respond or over-react to a non-threat than risk not reacting. For example, brushtail possums, *Trichosurus vulpecula*, respond more strongly to SPCs when there is no access to shelter and therefore greater vulnerability to predation (Parsons & Blumstein 2010). Encountering SPCs also led to changes in group movement patterns. However, in contrast to our predictions, meerkat groups reduced the distance travelled in the hour following an SPC encounter. Meerkats’ respond with greater intensity to SPCs in larger groups and when a greater proportion of the group is interacting (Driscoll *et al*. 2020). Thus, by the group recruiting and responding to the cue and increases in other defensive behaviours, this may reduce time used for moving, resulting in reduced movement in the period following a SPC encounter. Contrary to much of the literature (Monclús *et al*. 2005; Zidar & Løvlie 2012; Zöttl *et al*. 2013; Kuijper *et al*. 2014; Garvey *et al*. 2016; Kern *et al*. 2017; Tanis *et al*. 2018), we found no change in sentinel rate following a SPC encounter. It is nevertheless possible that there may have been changes in other vigilance types such as quadrupedal vigilance (scanning on four legs) as reported in Zöttl *et al*. (2013), which were not recorded in this study. In an area of greater risk as indicated by a SPC, it may be more beneficial for several individuals to perform scanning vigilance, rather than one sentinel being vigilant for the whole group. Increased scanning could also contribute to the reduction in distance travelled by, rather than reducing time for movement, slowing the movement itself with group members moving more cautiously and scanning.

Consistent with previous experimental findings (Driscoll *et al*. 2020), our results suggest that the mobbing-like response to SPCs does not act as a form of teaching. Thus, while meerkats are known to teach their pups to hunt (Thornton & McAuliffe 2006), there is no evidence they teach in any other context. In contrast to what would be predicted if SPC responses functioned to teach pups about potential threats, we found the frequency of SPC recruitment events was lower when pups were present and declined as the proportion of pups in the group increased. This is line with previous experimental work showing an inhibitory effect of pup presence on mobbing-like response intensity (decreased recruitment, tail raising and piloerection duration) (Driscoll *et al*. 2020). The influence of pup presence suggests the likelihood and strength of the mobbing-like response to SPCs is a plastic behaviour influenced by current constraints and conditions. We previously suggested the reduction in the intensity of SPC responses in the presence of pups may be the result of a trade-off between responding to SPCs and provisioning pups (Driscoll *et al*. 2020). In the current study, we found that the pup provisioning rate did not change following SPC encounters and was only constrained by the number of pups in the group, with the rate per pup declining as number of pups increased. Together, these findings suggest that when pups are present meerkats may reduce their investment in mobbing-like response to SPCs in order to maintain the pup provisioning rate. In contrast, there was no effect of the presence of pups on mobbing rate of live animals, suggesting lower plasticity in responding to actual predators with this response maintained irrespective of social conditions.

Many animals inspect SPCs (Brown & Godin 1999; Belton *et al*. 2007; Amo *et al*. 2011; Mella *et al*. 2014; Garvey *et al*. 2016), but recruitment to them is rare and, to our knowledge, has only been described in social mongoose species, with meerkats seemingly unique in displaying a mobbing-like response (Furrer & Manser 2009; Collier *et al*. 2017). This recruitment may facilitate collective-decision making in the face of threat uncertainty, rather than relying on the single encountering individual alarm calling making a decision on threat level and type alone. The recruitment of individuals may ensure the transfer of threat specific information, more so than may be obtainable from an alarm call, better informing the groups defensive behaviour. The exaggerated mobbing-like response may further increase likelihood of recruitment, by providing a conspicuous target for group members to recruit to. While there is scant evidence of recruitment to SPCs outside of mongoose species, this does not mean it does not occur. An analogous behaviour may be the inspection of SPCs in several fish species and the release of alarm cues to alert conspecifics to the threat (Brown & Godin 1999; Brown & Magnavacca 2003). However, it is not yet clear whether these alarm cues are used to only alert conspecifics to the threat, or also to initiate recruitment and inspection.

Overall the results indicate SPC encounters may be providing an indication of risk and informing group-level defensive behaviours. SPC encounters influenced subsequent behaviour, with an increase in alarm calling, indicative of increased risk, and a decrease in distance travelled, suggesting an increase in defensive behaviours. Also, the presence of pups affected the response to cues, with recruitment event frequency reducing when pups were present. The effects of pup presence and abiotic conditions implies there may be a certain degree of plasticity in responding to SPCs, more so than for mobbing of live animals. This plasticity may reflect condition dependent use and the necessity for using information gathered from SPCs in informing subsequent behaviour. Increasing our understanding of how animals respond to SPCs may provide important insights into the role of information use in shaping collective defensive behaviours.

## Acknowledgments

We thank the Kalahari Research Trust and the Northern Cape Conservation Authority for research permission (FAUNA 1020/2016). We also thank Dave Gaynor and Tim Vink for organising the field site, as well as the managers Coline Muller and Jacob Brown, and volunteers of the Kalahari Meerkat Project for organising, providing support and helping to collect the data and maintain habituation of the meerkats. Furthermore, we thank Michael Cant and Andrew Radford for providing valuable comments which helped improve this manuscript.

## Notes

### Competing Interest Statement

The authors have declared no competing interest.

