## Supplementary material for "Mobbing-like response to secondary predator cues informs group behaviour in wild meerkats"

***Recruitment event rate***

*Table 1. – Model summaries of the top candidate models using AIC model selection for predicting frequency of recruitment event to (a) secondary predator cues and (b) animals.*

| | Estimate | Std Error | ChiSq | df | p value |
|---|---|---|---|---|---|
| **a. Secondary predator cue** | | | | | |
| ***(a13) Proportion of pups* Average group size*** | | | | | |
| Intercept | 0.684 | 0.086 | | | |
| Prop. pups | -0.233 | 0.509 | 13.992 | 6 | <0.001 |
| Group size | 0.046 | 0.005 | 83.228 | 6 | <0.001 |
| Prop. pups:Group size | -0.064 | 0.037 | 10.750 | 7 | <0.001 |
| ***(a12) Pups foraging (y/n)* Average group size*** | | | | | |
| Intercept | 0.689 | 0.087 | | | |
| Forage | -0.287 | 0.149 | 22.789 | 6 | <0.001 |
| Group size | 0.046 | 0.005 | 93.796 | 6 | <0.001 |
| Forage:Group size | -0.001 | 0.008 | 0.013 | 7 | 0.911 |
| **b. Animal** | | | | | |
| ***(b6) Average group size*** | | | | | |
| Intercept | 0.311 | 0.052 | | | |
| Group size | 0.025 | 0.003 | 68.588 | 5 | <0.001 |
| ***(b13) Proportion* of pups** Average group size*** | | | | | |
| Intercept | 0.325 | 0.054 | | | |
| Prop. pups | -0.223 | 0.388 | 2.386 | 6 | 0.122 |
| Group size | 0.025 | 0.003 | 67.642 | 6 | <0.001 |
| Prop. pups:Group size | -0.004 | 0.028 | 0.413 | 7 | 0.520 |
| ***(b12) Pups foraging (y/n)* Average group size*** | | | | | |
| Intercept | 0.317 | 0.055 | | | |
| Forage | -0.044 | 0.113 | 0.634 | 6 | 0.426 |
| Group size | 0.025 | 0.003 | 69.217 | 6 | <0.001 |
| Forage:Group size | 0.000 | 0.006 | 0.010 | 7 | 0.921 |

### a. *Secondary predator cue rate*

*Table 2 – Model selection table for the factors affecting frequency of SPC recruitment events ranked by AIC value. Variables tested are pups foraging with the group (y/n), previous 30 day total rainfall, previous 9 month total rainfall, previous 30 days average maximum daily temperature, proportion of females in the group, proportion of pups in the group. Models forming the top set in bold.*

| | Intercept | Forage | Rainfall 30Days | Rainfall 9Months | MaxTemp 30Days | Prop F | Group Size | Prop Pups | Forage: Rainfall 30Days | Forage :Rainfall 9Months | Forage: MaxTemp 30Days | Forage: PropF | GroupSize: Forage | GroupSize: PropPups | MaxTemp3 0Days: Rainfall 9Months | MaxTemp 30Days: Rainfall 30Days | df | logLik | AICc | delta | weight |
|---|---|---|---|---|---|---|---|---|---|---|---|---|---|---|---|---|---|---|---|---|---|
| **a13** | **0.67** | | | | | | **0.05** | **0.07** | | | | | | **-0.08** | | | **7** | **-5025.62** | **10065.27** | **0.00** | **0.73** |
| **a12** | **0.69** | **+** | | | | | **0.05** | | | | | | **+** | | | | **7** | **-5026.59** | **10067.21** | **1.94** | **0.27** |
| **a6** | 0.66 | | | | | | 0.04 | | | | | | | | | | 5 | -5037.99 | 10086.00 | 20.72 | 0.00 |
| **a14** | -0.93 | | | 9.43 | 0.05 | | | | | | | | | | -0.23 | | 7 | -5040.63 | 10095.30 | 30.03 | 0.00 |
| **a9** | 0.87 | + | | 2.26 | | | | | | + | | | | | | | 7 | -5047.35 | 10108.74 | 43.47 | 0.00 |
| **a3** | 0.86 | | | 2.11 | | | | | | | | | | | | | 5 | -5054.12 | 10118.27 | 52.99 | 0.00 |
| **a15** | 1.74 | | 11.70 | | -0.02 | | | | | | | | | | | -0.24 | 7 | -5063.14 | 10140.32 | 75.05 | 0.00 |
| **a8** | 1.25 | + | 3.26 | | | | | | + | | | | | | | | 7 | -5065.24 | 10144.52 | 79.25 | 0.00 |
| **a11** | 1.61 | + | | | | -0.64 | | | | | | + | | | | | 7 | -5069.79 | 10153.63 | 88.36 | 0.00 |
| **a10** | 1.65 | + | | | -0.01 | | | | | | + | | | | | | 7 | -5070.47 | 10154.98 | 89.71 | 0.00 |
| **a7** | 1.30 | | | | | | | -0.70 | | | | | | | | | 5 | -5072.61 | 10155.23 | 89.96 | 0.00 |
| **a2** | 1.20 | | 2.77 | | | | | | | | | | | | | | 5 | -5073.32 | 10156.66 | 91.39 | 0.00 |
| **a1** | 1.31 | + | | | | | | | | | | | | | | | 5 | -5073.49 | 10157.00 | 91.73 | 0.00 |
| **a4** | 1.66 | | | | -0.01 | | | | | | | | | | | | 5 | -5076.18 | 10162.38 | 97.11 | 0.00 |
| **a5** | 1.50 | | | | | -0.51 | | | | | | | | | | | 5 | -5078.06 | 10166.13 | 100.86 | 0.00 |

### b. *Animal mobbing rate*

*Table 3 – Model selection table for the factors affecting frequency of animal recruitment events ranked by AIC value. Variables tested are pups foraging with the group (y/n), previous 30 day total rainfall, previous 9 month total rainfall, previous 30 days average maximum daily temperature, proportion of females in the group, proportion of pups in the group. Models forming the top set in bold.*

| | Intercept | Forage | Rainfall 30DaysL | Rainfall 9Months | MaxTemp 30Days | Prop F | Group size | PropPups | Forage:Rainfall 30Days | Forage:Rainfall 9Months | Forage:MaxTemp 30Days | Forage:Prop F | Group Size:Forage | GroupSize:Prop pups | MaxTemp 30Days:Rainfall 9Months | MaxTemp 30Days:Rainfall 30Days | df | logLik | AICc | delta | weight |
|---|---|---|---|---|---|---|---|---|---|---|---|---|---|---|---|---|---|---|---|---|---|
| **b6** | **0.31** | | | | | | **0.03** | | | | | | | | | | **5** | **-4235.79** | **8481.61** | **0.00** | **0.58** |
| **b13** | **0.32** | | | | | | **0.03** | **-0.10** | | | | | | **-0.01** | | | **7** | **-4234.39** | **8482.82** | **1.22** | **0.31** |
| **b12** | **0.32** | **+** | | | | | **0.03** | | | | | | **+** | | | | **7** | **-4235.47** | **8484.98** | **3.37** | **0.11** |
| **b4** | 0.27 | | | | 0.01 | | | | | | | | | | | | 5 | -4262.32 | 8534.66 | 53.05 | 0.00 |
| **b14** | -0.21 | | | 2.10 | 0.03 | | | | | | | | | | -0.06 | | 7 | -4260.51 | 8535.05 | 53.45 | 0.00 |
| **b10** | 0.34 | + | | | 0.01 | | | | | | + | | | | | | 7 | -4261.09 | 8536.23 | 54.62 | 0.00 |
| **b15** | 0.34 | | -3.56 | | 0.01 | | | | | | | | | | | 0.13 | 7 | -4261.22 | 8536.48 | 54.87 | 0.00 |
| **b2** | 0.65 | | 1.43 | | | | | | | | | | | | | | 5 | -4266.93 | 8543.89 | 62.28 | 0.00 |
| **b8** | 0.64 | + | 1.87 | | | | | | + | | | | | | | | 7 | -4266.09 | 8546.22 | 64.62 | 0.00 |
| **b7** | 0.69 | | | | | | | -0.25 | | | | | | | | | 5 | -4268.42 | 8546.86 | 65.26 | 0.00 |
| **b5** | 0.82 | | | | | -0.30 | | | | | | | | | | | 5 | -4268.54 | 8547.10 | 65.49 | 0.00 |
| **b3** | 0.74 | | | -0.33 | | | | | | | | | | | | | 5 | -4268.88 | 8547.77 | 66.17 | 0.00 |
| **b11** | 0.79 | + | | | | -0.23 | | | | | | + | | | | | 7 | -4267.96 | 8549.96 | 68.35 | 0.00 |
| **b9** | 0.72 | + | | -0.20 | | | | | | + | | | | | | | 7 | -4267.97 | 8549.99 | 68.38 | 0.00 |
| **b1** | 0.68 | + | | | | | | | | | | | | | | | 5 | -4270.08 | 8550.19 | 68.58 | 0.00 |

***Behavioural changes following recruitment event***

*Table 4. – Model summaries of the top candidate models following AIC model selection for various behaviours that may change following a SPC recruitment event; (c) alarm calling rate, (d) guarding rate, (f) distance travelled, (e) per pup provisioning rate.*

| | Estimate | Std Error | ChiSq | df | p value |
|---|---|---|---|---|---|
| **c. Alarm calling rate** | | | | | |
| ***(c8) Hour before or after*Daily maximum temperature*** | | | | | |
| Intercept | 1.247 | 0.084 | | | |
| Before/after | 0.033 | 0.110 | 13.344 | 6 | <0.001 |
| Max. Temp. | -0.015 | 0.003 | 73.606 | 6 | <0.001 |
| Before or after:Max. temp. | -0.004 | 0.004 | 1.225 | 7 | 0.269 |
| ***(c12) Daily maximum temperature*Rainfall 9 Months*** | | | | | |
| Intercept | 1.567 | 0.162 | | | |
| Max. Temp. | -0.028 | 0.005 | 46.398 | 6 | <0.001 |
| Rainfall 9 Months | -1.432 | 0.582 | 0.903 | 6 | 0.342 |
| Max. Temp.*Rainfall 9 Months | 0.054 | 0.019 | 8.086 | 7 | 0.004 |
| **d. Sentinel rate** | | | | | |
| ***(d19) Pups foraging (y/n)*Rainfall 9 Months*** | | | | | |
| Intercept | -0.200 | 0.071 | | | |
| Forage | 0.351 | 0.087 | 16.297 | 6 | <0.001 |
| Rainfall 9 Months | 2.748 | 0.206 | 171.230 | 6 | <0.001 |
| Forage*Rainfall 9 Months | -0.847 | 0.331 | 6.505 | 7 | 0.011 |
| **e. Distance travelled** | | | | | |
| ***(e8) Hour before or after*Max. temp.*** | | | | | |
| Intercept | 4.481 | 0.087 | | | |
| Before/after | 0.268 | 0.116 | 18.230 | 6 | <0.001 |
| Max. temp. | 0.019 | 0.003 | 68.113 | 6 | <0.001 |
| Before/after *Max. temp. | -0.005 | 0.004 | 2.115 | 7 | 0.146 |
| ***(e12) Daily maximum temperature*Rainfall 9 Months*** | | | | | |
| Intercept | 4.668 | 0.164 | | | |
| Max. temp. | 0.019 | 0.005 | 26.982 | 6 | <0.001 |
| Rainfall 9 Months | 0.343 | 0.658 | 13.897 | 6 | <0.001 |
| Max. temp.*Rainfall 9 Months | -0.033 | 0.021 | 2.472 | 7 | 0.116 |
| ***(e13) Daily maximum temperature *Rainfall 30 days*** | | | | | |
| Intercept | 4.526 | 0.075 | | | |

| | | | | | |
|---|---|---|---|---|---|
| Max. Temp. | 0.020 | 0.002 | 75.939 | 6 | <0.001 |
| Rainfall 30 days | 4.079 | 2.608 | 11.706 | 6 | 0.001 |
| Max. temp.*Rainfall 30 Days | -0.170 | 0.080 | 4.484 | 7 | 0.034 |
| **Per pup provisioning rate** | | | | | |
| ***(f14) Proportion of pups*** | | | | | |
| Intercept | 0.322 | 0.233 | | | |
| Prop. pups | -2.864 | 0.626 | 22.528 | 6 | <0.001 |
| ***(f15) Proportion of pups*Average group size*** | | | | | |
| Intercept | 0.160 | 0.458 | | | |
| Prop. pups | -0.793 | 1.441 | 21.378 | 7 | <0.001 |
| Group size | 0.013 | 0.018 | 1.246 | 7 | 0.264 |
| Prop. pups*Group size | -0.147 | 0.076 | 3.611 | 8 | 0.057 |
| ***(f3) Proportion of pups*Hour before or after*** | | | | | |
| Intercept | 0.289 | 0.248 | | | |
| Prop. pups | -2.785 | 0.729 | 22.468 | 7 | <0.001 |
| Before/after | 0.065 | 0.170 | 0.233 | 7 | 0.629 |
| Prop. pups* Before/after | -0.154 | 0.762 | 0.041 | 8 | 0.840 |

#### *c. Alarm calling rate*

*Table 5 – Model selection table for the factors affecting hourly alarm calling rate ranked by AIC value. Variables tested are the hour before or after a recruitment event, pups foraging with the group (y/n), proportion of pups in the group, predator cue type (scent or object), previous 30 day total rainfall, previous 9 month total rainfall, daily maximum temperature. Models forming the top set in bold.*

| Model | Intercept | B/A | Forage | B/A:Forage | Pred Code | B/A:PredCode | PropPups | B/A:PropPups | GroupSize | B/A:GroupSize | Rainfall9MonthsL | B/A:Rainfall 9Months | Rainfall30Days | AlarmFac:Rainfall [illegible] | MaxTemp | B/A:MaxTemp | MaxTemp:Rainfall9Months | MaxTemp:Rainfall 30Days | GroupSize:PropPups | Forage:PredCode | Forage:MaxTemp | Forage:Rainfall 9Months | Forage:Rainfall 30Days | Forage:GroupSize | df | logLik | AIC | delta | weight |
|---|---|---|---|---|---|---|---|---|---|---|---|---|---|---|---|---|---|---|---|---|---|---|---|---|---|---|---|---|---|
| **c8** | **1.25** | **+** | | | | | | | | | | | | | **-0.01** | **+** | | | | | | | | | **7** | **-13605.14** | **27224.28** | **0.00** | **0.92** |
| **c12** | **1.57** | | | | | | | | | | **-1.43** | | | | **-0.03** | | **0.05** | | | | | | | | **7** | **-13607.93** | **27229.86** | **5.58** | **0.06** |
| **c13** | 1.35 | | | | | | | | | | | | -4.71 | | -0.02 | | | 0.16 | | | | | | | 7 | -13609.17 | 27232.34 | 8.06 | 0.02 |
| **c22** | 1.26 | | | | | | | | | | | | | | -0.02 | | | | | | | | | | 5 | -13612.42 | 27234.85 | 10.57 | 0.00 |
| **c18** | 1.22 | | + | | | | | | | | | | | | -0.02 | | | | | | + | | | | 7 | -13611.08 | 27236.16 | 11.88 | 0.00 |
| **c6** | 0.62 | + | | | | | | | | | 0.84 | + | | | | | | | | | | | | | 7 | -13628.43 | 27270.87 | 46.59 | 0.00 |
| **c11** | 0.60 | | | | | | | | | | 0.74 | | | | | | | | | | | | | | 5 | -13635.17 | 27280.34 | 56.06 | 0.00 |
| **c19** | 0.62 | | + | | | | | | | | 0.72 | | | | | | | | | | | + | | | 7 | -13633.58 | 27281.16 | 56.88 | 0.00 |
| **c2** | 0.81 | + | + | + | | | | | | | | | | | | | | | | | | | | | 7 | -13639.13 | 27292.26 | 67.98 | 0.00 |
| **c4** | 0.79 | + | | | | | 0.38 | + | | | | | | | | | | | | | | | | | 7 | -13639.20 | 27292.40 | 68.12 | 0.00 |

| | | | | | | | | | | | | | | | | | | |
|---|---|---|---|---|---|---|---|---|---|---|---|---|---|---|---|---|---|---|
| **c1** | 0.81 | + | | | | | | | | | | | | | 5 | -13642.55 | 27295.11 | 70.83 | 0.00 |
| **c5** | 0.77 | + | | | | | 0.00 | + | | | | | | | 7 | -13641.15 | 27296.30 | 72.02 | 0.00 |
| **c7** | 0.81 | + | | | | | | | -0.26 | + | | | | | 7 | -13641.28 | 27296.57 | 72.29 | 0.00 |
| **c3** | 0.79 | + | | + | + | | | | | | | | | | 7 | -13642.50 | 27298.99 | 74.71 | 0.00 |
| **c16** | 0.74 | | | | | 1.00 | 0.00 | | | | -0.06 | | | | 7 | -13644.65 | 27303.29 | 79.01 | 0.00 |
| **c14** | 0.78 | | + | | | | | | | | | | | | 5 | -13647.18 | 27304.35 | 80.07 | 0.00 |
| **c21** | 0.76 | | + | | | | 0.00 | | | | | | | + | 7 | -13645.93 | 27305.86 | 81.58 | 0.00 |
| **c23** | 0.78 | | | | | | | | -0.48 | | | | | | 5 | -13647.95 | 27305.90 | 81.62 | 0.00 |
| **c20** | 0.79 | | + | | | | | | -0.49 | | | | + | | 7 | -13646.38 | 27306.77 | 82.49 | 0.00 |
| **c15** | 0.76 | | | | | 0.08 | | | | | | | | | 5 | -13648.77 | 27307.55 | 83.27 | 0.00 |
| **c10** | 0.77 | | | | | | 0.00 | | | | | | | | 5 | -13648.90 | 27307.80 | 83.53 | 0.00 |
| **c9** | 0.76 | | | + | | | | | | | | | | | 5 | -13648.90 | 27307.81 | 83.53 | 0.00 |
| **c17** | 0.78 | | + | + | | | | | | | | + | | | 7 | -13647.09 | 27308.18 | 83.90 | 0.00 |

*d. **Guarding rate***

*Table 6 – Model selection table for the factors affecting hourly guarding bout rate ranked by AIC value. Variables tested are the hour before or after a recruitment event, pups foraging with the group (y/n), proportion of pups in the group, predator cue type (scent or object), previous 30 day total rainfall, previous 9 month total rainfall, daily maximum temperature. Models forming the top set in bold.*

| | Intercept | B/A | Forage | Forage: B/A | PredCode | B/A:Pred Code | PropPups | B/A:Prop Pups | Group Size | Group Size: B/A | Rainfall 9Months | B/A:Rainfall9Months | Rainfall3 0Days | B/A:Rainfall30Days | Max Temp | B/A:Max Temp | MaxTemp1Day:Rainfall9 Months | Max TempRainfall 30Days | Group Size:Prop Pups | Forage:PredCode | Forage:MaxTemp1 | Forage:Rainfall 9Months | Forage:Rainfall 30Days | Forage:GroupSize | df | logLik | AICc | delta | weight |
|---|---|---|---|---|---|---|---|---|---|---|---|---|---|---|---|---|---|---|---|---|---|---|---|---|---|---|---|---|---|
| **d19** | **-0.20** | | **+** | | | | | | | | **2.75** | | | | | | | | | | | **+** | | | **7** | **-12248.16** | **24510.33** | **0.00** | **1.00** |
| **d6** | -0.18 | + | | | | | | | | | 2.82 | + | | | | | | | | | | | | | 7 | -12256.58 | 24527.18 | 16.85 | 0.00 |
| **d11** | -0.10 | | | | | | | | | | 2.51 | | | | | | | | | | | | | | 5 | -12259.56 | 24529.12 | 18.79 | 0.00 |
| **d12** | 0.17 | | | | | | | | | | 1.37 | | | | -0.01 | | 0.04 | | | | | | | | 7 | -12258.26 | 24530.54 | 20.21 | 0.00 |
| **d18** | 0.97 | | + | | | | | | | | | | | | -0.02 | | | | | | + | | | | 7 | -12310.32 | 24634.66 | 124.33 | 0.00 |
| **d22** | 0.97 | | | | | | | | | | | | | | -0.02 | | | | | | | | | | 5 | -12319.70 | 24649.40 | 139.08 | 0.00 |
| **d13** | 1.00 | | | | | | | | | | | | -0.34 | | -0.02 | | | 0.03 | | | | | | | 7 | -12318.98 | 24651.97 | 141.64 | 0.00 |
| **d8** | 0.93 | + | | | | | | | | | | | | | -0.02 | + | | | | | | | | | 7 | -12319.52 | 24653.06 | 142.74 | 0.00 |
| **d16** | 0.17 | | | | | | 0.94 | | 0.01 | | | | | | | | | | 0.00 | | | | | | 7 | -12322.14 | 24658.30 | 147.97 | 0.00 |
| **d15** | 0.40 | | | | | | 0.95 | | | | | | | | | | | | | | | | | | 5 | -12329.35 | 24668.71 | 158.39 | 0.00 |

| | | | | | | | | | | | | | | | | | | | | |
|---|---|---|---|---|---|---|---|---|---|---|---|---|---|---|---|---|---|---|---|---|
| **d21** | 0.14 | | + | | | | | | 0.02 | | | | | | + | 7 | -12328.55 | 24671.12 | 160.79 | 0.00 |
| **d4** | 0.38 | + | | | | | 1.09 | + | | | | | | | | 7 | -12328.91 | 24671.83 | 161.50 | 0.00 |
| **d5** | 0.10 | + | | | | | | | 0.02 | + | | | | | | 7 | -12330.86 | 24675.73 | 165.40 | 0.00 |
| **d10** | 0.20 | | | | | | | | 0.01 | | | | | | | 5 | -12334.44 | 24678.90 | 168.57 | 0.00 |
| **d17** | 0.27 | | + | | + | | | | | | | | + | | | 7 | -12333.63 | 24681.28 | 170.95 | 0.00 |
| **d14** | 0.41 | | + | | | | | | | | | | | | | 5 | -12337.02 | 24684.05 | 173.73 | 0.00 |
| **d20** | 0.41 | | + | | | | | | | | -0.25 | | | + | | 7 | -12336.42 | 24686.85 | 176.52 | 0.00 |
| **d2** | 0.39 | + | + | + | | | | | | | | | | | | 7 | -12336.60 | 24687.22 | 176.90 | 0.00 |
| **d9** | 0.33 | | | | + | | | | | | | | | | | 5 | -12339.81 | 24689.62 | 179.29 | 0.00 |
| **d3** | 0.34 | + | | | + | + | | | | | | | | | | 7 | -12339.70 | 24693.41 | 183.09 | 0.00 |
| **d23** | 0.45 | | | | | | | | | | -0.28 | | | | | 5 | -12342.43 | 24694.86 | 184.53 | 0.00 |
| **d1** | 0.44 | + | | | | | | | | | | | | | | 5 | -12342.59 | 24695.19 | 184.86 | 0.00 |
| **d7** | 0.45 | + | | | | | | | | | -0.50 | + | | | | 7 | -12342.22 | 24698.46 | 188.14 | 0.00 |

### *e. Distance travelled*

*Table 7 – Model selection table for the factors distance travelled in an hour ranked by AIC value. Variables tested are the hour before or after a recruitment event, pups foraging with the group (y/n), proportion of pups in the group, predator cue type (scent or object), previous 30 day total rainfall, previous 9 month total rainfall, daily maximum temperature. Models forming the top set in bold.*

| | Intercept | B/A | Forage | B/A:Forage | PredCode | B/A:PredCode | Prop Pups | B/A:Prop Pups | Group Size | B/A:Group Size | Rainfall 9Months | B/A:Rainfall 9Months | Rainfall 30Days | B/A:Rainfall 30Days | Max Temp | B/A:Max Temp | Max Temp:Rainfall 9Months | Max Temp:Rainfall 30Days | Group Size:Prop Pups | Forage:Pred Code | Forage:Max Temp | Forage:Rainfall 9Months | Forage:Rainfall3 0Days | Forage:Group Size | df | logLik | AICc | delta | weight |
|---|---|---|---|---|---|---|---|---|---|---|---|---|---|---|---|---|---|---|---|---|---|---|---|---|---|---|---|---|---|
| **e8** | **4.48** | **+** | | | | | | | | | | | | | **0.02** | **+** | | | | | | | | | **7** | **-24463.53** | **48941.09** | **0.00** | **0.79** |
| **e12** | **4.67** | | | | | | | | | | **0.34** | | | | **0.02** | | **-0.03** | | | | | | | | **7** | **-24465.52** | **48945.07** | **3.98** | **0.11** |
| **e13** | **4.53** | | | | | | | | | | | | **4.08** | | **0.02** | | | **-0.17** | | | | | | | **7** | **-24465.61** | **48945.24** | **4.15** | **0.10** |
| **e6** | 5.30 | + | | | | | | | | | -1.17 | + | | | | | | | | | | | | | 7 | -24471.49 | 48957.00 | 15.91 | 0.00 |
| **e22** | 4.62 | | | | | | | | | | | | | | 0.02 | | | | | | | | | | 5 | -24473.70 | 48957.42 | 16.33 | 0.00 |
| **e18** | 4.64 | | + | | | | | | | | | | | | 0.02 | | | | | | + | | | | 7 | -24472.60 | 48959.24 | 18.15 | 0.00 |
| **e11** | 5.34 | | | | | | | | | | -1.11 | | | | | | | | | | | | | | 5 | -24480.25 | 48970.51 | 29.42 | 0.00 |
| **e19** | 5.34 | | + | | | | | | | | -1.08 | | | | | | | | | | | + | | | 7 | -24480.01 | 48974.04 | 32.95 | 0.00 |
| **e4** | 5.09 | + | | | | | -0.24 | + | | | | | | | | | | | | | | | | | 7 | -24495.75 | 49005.52 | 64.43 | 0.00 |
| **e1** | 5.08 | + | | | | | | | | | | | | | | | | | | | | | | | 5 | -24498.64 | 49007.30 | 66.21 | 0.00 |

| | | | | | | | | | | | | | | | | | | | | |
|---|---|---|---|---|---|---|---|---|---|---|---|---|---|---|---|---|---|---|---|---|
| **e7** | 5.10 | + | | | | | | | | -0.80 | + | | | | | 7 | -24497.17 | 49008.38 | 67.29 | 0.00 |
| **e5** | 5.14 | + | | | | | | 0.00 | + | | | | | | | 7 | -24497.35 | 49008.73 | 67.64 | 0.00 |
| **e3** | 5.02 | + | | | + | + | | | | | | | | | | 7 | -24497.87 | 49009.76 | 68.67 | 0.00 |
| **e2** | 5.09 | + | + | + | | | | | | | | | | | | 7 | -24498.27 | 49010.56 | 69.47 | 0.00 |
| **e15** | 5.14 | | | | | | -0.31 | | | | | | | | | 5 | -24504.42 | 49018.85 | 77.76 | 0.00 |
| **e23** | 5.14 | | | | | | | | | -0.66 | | | | | | 5 | -24505.82 | 49021.65 | 80.56 | 0.00 |
| **e16** | 5.14 | | | | | | -0.05 | 0.00 | | | | -0.02 | | | | 7 | -24503.96 | 49021.94 | 80.85 | 0.00 |
| **e9** | 5.09 | | | | + | | | | | | | | | | | 5 | -24506.55 | 49023.11 | 82.02 | 0.00 |
| **e14** | 5.13 | | + | | | | | | | | | | | | | 5 | -24507.10 | 49024.22 | 83.13 | 0.00 |
| **e10** | 5.15 | | | | | | | 0.00 | | | | | | | | 5 | -24507.13 | 49024.27 | 83.18 | 0.00 |
| **e20** | 5.15 | | + | | | | | | | -0.69 | | | | + | | 7 | -24505.78 | 49025.59 | 84.50 | 0.00 |
| **e17** | 5.07 | | + | | + | | | | | | | | + | | | 7 | -24505.99 | 49026.01 | 84.92 | 0.00 |
| **e21** | 5.14 | | + | | | | | 0.00 | | | | | | | + | 7 | -24506.92 | 49027.86 | 86.77 | 0.00 |

### *f. Pup provisioning*

*Table 8. – Model selection table for the factors affecting per pup provisioning rate ranked by AIC value. Variables tested are the hour before or after a recruitment event, proportion of pups in the group, predator cue type (scent or object), previous 30 day total rainfall, previous 9 month total rainfall, daily maximum temperature. Models forming the top set in bold.*

| | Intercept | ZI Intercept | B/A | PredCode | Pred Code: B/A | PropPups | PropPups: B/A | GroupSize | Group Size: B/A | Rainfall9MonthsL | B/A:Rainfall 9Months | Rainfall30DaysL | B/A:Rainfall 30Days | Max Temp | Max Temp: B/A | MaxTemp:Rainfall9Months | Max Temp:Rainfall 30Days | GroupSize:PropPups | df | logLik | AICc | delta | weight |
|---|---|---|---|---|---|---|---|---|---|---|---|---|---|---|---|---|---|---|---|---|---|---|---|
| **f14** | **0.16** | **-21.06** | | | | **-0.79** | | **0.01** | | | | | | | | | | **-0.15** | **8** | **-1807.13** | **3630.38** | **0.00** | **0.57** |
| **f13** | **0.32** | **-20.77** | | | | **-2.86** | | | | | | | | | | | | | **6** | **-1809.56** | **3631.19** | **0.81** | **0.38** |
| **f3** | **0.29** | **-21.16** | **+** | | | **-2.78** | **+** | | | | | | | | | | | | **8** | **-1809.42** | **3634.96** | **4.58** | **0.06** |
| **f2** | -0.14 | -21.32 | + | + | + | | | | | | | | | | | | | | 8 | -1816.40 | 3648.92 | 18.54 | 0.00 |
| **f10** | -0.11 | -20.76 | | | | | | | | -0.90 | | | | | | | | | 6 | -1819.40 | 3650.86 | 20.48 | 0.00 |
| **f9** | -0.59 | -21.07 | | | | | | 0.01 | | | | | | | | | | | 6 | -1819.63 | 3651.32 | 20.94 | 0.00 |
| **f8** | -0.46 | -20.93 | | + | | | | | | | | | | | | | | | 6 | -1820.31 | 3652.69 | 22.31 | 0.00 |
| **f11** | 0.01 | -21.34 | | | | | | | | -0.19 | | | | 0.00 | | -0.03 | | | 8 | -1818.59 | 3653.30 | 22.92 | 0.00 |
| **f1** | -0.33 | -21.19 | + | | | | | | | | | | | | | | | | 6 | -1820.68 | 3653.42 | 23.04 | 0.00 |
| **f16** | -0.29 | -20.88 | | | | | | | | | | -0.50 | | | | | | | 6 | -1820.71 | 3653.49 | 23.11 | 0.00 |
| **f15** | -0.24 | -20.87 | | | | | | | | | | | | 0.00 | | | | | 6 | -1820.77 | 3653.62 | 23.24 | 0.00 |

| | | | | | | | | | | | | | | | | | |
|---|---|---|---|---|---|---|---|---|---|---|---|---|---|---|---|---|---|
| **f5** | -0.21 | -21.30 | + | | | -0.55 | + | | | | | | 8 | -1818.78 | 3653.68 | 23.30 | 0.00 |
| **f4** | -0.65 | -20.84 | + | 0.02 | + | | | | | | | | 8 | -1819.37 | 3654.86 | 24.47 | 0.00 |
| **f12** | -0.06 | -13.21 | | | | | | -10.80 | | -0.01 | | 0.31 | 8 | -1819.86 | 3655.84 | 25.46 | 0.00 |
| **f7** | -0.38 | -20.89 | + | | | | | | | 0.00 | + | | 8 | -1820.45 | 3657.02 | 26.64 | 0.00 |
| **f6** | -0.32 | -20.34 | + | | | | | -0.35 | + | | | | 8 | -1820.55 | 3657.22 | 26.84 | 0.00 |
